# Lytic *Bacteroides uniformis* bacteriophages exhibiting host tropism congruent with diversity generating retroelement

**DOI:** 10.1101/2020.10.09.334284

**Authors:** Stina Hedzet, Tomaž Accetto, Maja Rupnik

**Affiliations:** National laboratory for health, environment and food, NLZOH, Maribor, Slovenia; University of Ljubljana, Biotechnical faculty, Animal science department; University of Maribor, Faculty of Medicine, Maribor, Slovenia

**Author notes:** Corresponding author: Maja Rupnik, NLZOH.

**Keywords:** gut, *Bacteroides*, virome, lytic phage, prophage, Diversity Generating Retroelement

## Abstract

Intestinal phages are abundant and important component of gut microbiota, but our knowledge remains limited to only a few isolated and characterized representatives targeting numerically dominant gut bacteria. Here we describe isolation of human intestinal phages infecting *Bacteroides uniformis. Bacteroides* is one of the most common bacterial groups in the global human gut microbiota, however, to date not many *Bacteroides* specific phages are known. Phages isolated in this study belong to a novel viral genus, Bacuni, within *Siphoviridae* family and represent the first lytic phages, genomes of which encode diversity generating retroelements (DGR). This region is assumed to promote phage adaptation to the rapidly changing environmental conditions and to broaden its host range. Three isolated phages showed 99,83% genome identity but infected distinct *B. uniformis* strains. The tropism of Bacuni phages appeared to be dependent on the interplay of DGR mediated sequence variations of phage fimbrial tip proteins and mutations in host genes coding for outer-membrane proteins. We found prophages with up to 85% aa similarity to Bacuni phages in the genomes of *B. acidifaciens* and *Prevotella* sp.. Despite the abundance of *Bacteroides* within human microbiome, we found Bacuni phages only in a limited subset of published gut metagenomes.

**Importance:** The lack of common marker gene in viruses require a precise characterization of diverse isolated phages to enhance metagenomic analyses and to understand their role in gut microbiota. Here we report the isolation of phages representing a new genus with characteristics so far not known or rarely described in intestinal phages. They are the first lytic phages specific for *Bacteroides uniformis*, a bacterial representative of the prevalent genus in the gut of humans and animals. Additionally, they are the first lytic phages containing specific regions (diversity generating retroelement) that putatively influence host tropism. The ability to switch constantly the targeted populations of the host species could provide an evolutionary advantage to these bacteriophages and may affect intra species diversity.

## Introduction

Intestinal viruses and their impact on human health are a neglected component of the widely studied gut microbiota. Bacteriophages (phages) exhibit different life styles and play an important role in shaping bacterial diversity and composition of the intestinal microbiota through predation and horizontal gene transfer ^1,2^. Sequencing-based metagenomic studies have enabled insight into this complex viral reservoir revealing genetically very diverse phages ^1,3–5^.

Virome metagenomic studies encounter several difficulties. The vast majority (75-99%) of sequencing reads does not correspond to any matches in the existing viral databases ^3^. Viruses lack universal marker genes, while standardized protocols for sample preparations and analysis are not yet established ^1^. To decipher gut virome and to connect biological characteristics with metagenomic data, cultivation of intestinal phages and their associated hosts remains crucial. A great number of intestinal phages infect anaerobic bacteria, which are challenging to cultivate; isolated and characterized phages are therefore sparse.

Despite these difficulties, several phages and prophages were lately described in different anaerobic gut microbiota representatives. *In silico* discovered viral clade, CrAss-like phages, is presumably present in 50% of Western individuals and can represent up to 90% of viral metagenomics reads per individual sample ^6,7^. Prediction of suspected *Bacteroides* sp. host was confirmed by isolation of a CrAss-like phage, Crass001, that infects *Bacteroides intestinalis* and exhibits a podovirus-like morphology ^8^. Its life style has yet to be elucidated. CrAss-like phages are a group of genetically highly diverse phages making additional *Bacteroides* sp. or other bacteria probable hosts ^9^. Lysogenic (temperate) phages have been identified in genomes of *Fecalibacterium prausnitzii* ^10^ and *Bacteroides dorei* ^11^. Extremely large gut phage genomes (540 kb), named Lak phages, that presumably infect *Prevotella sp*. were also recovered from gut metagenomes ^12^. Recently, a study of temperate phage–bacteria interactions in mice gut showed that *Roseburia intestinalis* prophages influence temporal variations in composition of gut microbiota ^13^. Additionally to *Bacteroides dorei* Hankyphage ^11^ and CrAss001 ^8^, four phages infecting different species within *Bacteroides* genus were isolated and sequenced. Phages B40-8 ^14^ and B124-14 ^15^ infect *Bacteroides fragilis*, while phages ϕBrb01 and ϕBrb02, originating from sewage, infect *Bacteroides* sp. bacterial hosts isolated from rumen fluid ^16^. However, compared to more than 150 phages infecting *E. coli* isolated from various biomes and clinical settings ^17^, there are few reported bacteriophages infecting species from the genus *Bacteroides*, which account for roughly 30% of all bacteria in an average human intestine ^18^.

Diversity-generating retroelements (DGRs) are genetic elements, composed of template-dependent reverse transcriptase and accessory proteins that produce mutations in targeted genes with variable repeats. This introduces the variability in the target proteins ^19^. DGR mechanism was first described in *Bordetella* phage BPP-1, in which mutations target phage tail fiber gene via “mutagenic retrohoming” to enable bacterial host species switching ^20,21^. Phage-encoded DGRs were also found in genomes of isolated temperate phages of intestinal *B. dorei* ^11^ and *F. prausnitzii* ^10^. Moreover, DGRs were detected in defined prophage regions of bacteria belonging to *Bacteroidetes, Proteobacteria* and *Firmicutes*, obtained from human-gut associated metagenomes and bacterial genomes ^11^.

Understanding the intestinal virome depends on the number of isolated, sequenced, and characterized bacteriophages and their associated hosts. The aim of the present study was to obtain and characterize the phages targeting abundant gut bacteria from *Bacteroides* genus, and to contribute to the insight of the “viral dark matter” ^22^ of the interactions of bacteria and viruses in human gut. Additionally, the study provides bioinformatic evidence that the host range of isolated phages may very well be mediated by a DGR.

## Materials and methods

### Isolation of bacterial strains from human fecal sample

Fecal sample, obtained from a healthy volunteer was aliquoted and further processed or stored at −80°C. The complete isolation of bacterial strains and preparation of fecal suspension was carried out in an anaerobic workstation at 37 °C (Don Whitley Scientific).

Dilutions of homogenized fecal suspension (20%), made from fresh feces and pre-reduced anaerobic YBHI culture media (Brain-heart infusion media, supplemented with 0.5 % yeast extract (BioLife) and 20% of rumen fluid) were plated on YBHI agar. After 72 hours of anaerobic incubation at 37°C single colonies were randomly chosen and isolated on YBHI plates to obtain pure bacterial cultures. Isolates were identified by mass spectrometry (MALDI-TOF Biotyper System, Bruker Daltonik, Bremen, Germany). Identification of *Bacteroides* strains was confirmed by 16S rRNA gene sequencing amplified with primers 27feb to 1495revb ^23^ and analyzed with RDP Classifier ^24^.

Isolated *Bacteroides* strains (Data set S1) were then used in phage screening experiment and host range experiment.

### Phage enrichment from sterile filtrate of homogenized fecal sample

Fecal sample used for the phage isolation was not identical as used for bacterial strain isolation but was retrieved from the same healthy volunteer. Fresh fecal material (5g) was resuspended in 50 mL of SM buffer with vigorous vortexing for 20 minutes. SM buffer contained 100 mM NaCl, 8 mM MgSO_4_, 50 mM Tris-Cl (1 M, pH 7.5) and 0.01% (w/v) gelatine (2%, w/v)). After cooling down on ice, fecal suspension was centrifuged twice at 5400 × g (4°C). Supernatant was filtered twice through 0.2 µm pore cellulose acetate (CA) syringe membrane filters (Filtropur, Starsted). Sterile filtrate of homogenized fecal sample (fecal water) was stored at +4°C until further use.

Phages were initially enriched in *Bacteroides* cultures. Ten different *Bacteroides* strains in stationary phase (1mL) were subcultured into 9 mL of liquid sABB (Anaerobe Basal Broth, Thermo Fisher Scientific, supplemented with MgSO_4_ (0.12 mM) and CaCl_2_ (1 mM)). For phage enrichment, 1 mL of fecal water was added to the inoculated media and incubated for 24 hours at 37°C. Subsequently, 3 mL of culture media were removed and centrifuged at 5400 × g (4°C). Supernatant was syringe-filtered (0.2 µm pore, Starsted) and added to 9 mL of fresh sABB media, inoculated with *Bacteroides* strain in stationary phase like described before. The procedure was again repeated after 24 hours. The final sterile supernatant was refrigerated (4°C) until further used in double-agar-layer method. Maximal storage duration was 72 hours.

### Phage isolation from enrichment co-cultures with Bacteroides

Spot assay on a double-agar-layer (DAL) was used for phage isolation from enrichment cultures.

*Bacteroides* strains, cultivated in liquid sABB, were sampled at two different time points with optical densities OD_620_ 0,2 (T1) and OD_620_ 0,5 (T2) for further use in DAL assay. For each time point 10-fold dilutions were made and 200 µL of each dilution was mixed with 3 ml soft agar that was kept anaerobically at 47°C (sABB) and poured on the prereduced sABB agar basal plates. After solidification 10-fold dilutions of supernatant filtrates of phage enrichment cultures (10µL) were spotted on solidified agar. After 24 h of incubation plates were checked for potential lysis zones. The top agar with clear zones was harvested with an inoculation loop and stored in 100 µL of SM puffer for 18-24 h at 4°C, followed by centrifugation (13 000 × g, 5 min). Supernatant was then used for further steps in phage purification and characterization.

Phages were purified from the stored spot assay supernatants by three consecutive single plaque isolation cycles using the corresponding bacterial host strain. *Bacteroides* culture (200 µL) in *log* growth phase was mixed with 10-fold dilutions of lysis zone supernatant and 2.5 mL of sABB soft agar and poured onto sABB agar basal plates, allowed to solidify, and incubated at 37°C. After 18-24h incubation, a single plaque was picked with pipette tip, transferred to SM buffer (100 µL) and left overnight at 4°C. After 18-24 h, phage lysates were centrifuged (13 000 × g, 5 min) and used in a plaque assay.

### Preparation of phage stock suspensions, EM characterization and host range

Each isolated phage in SM buffer (100 µL) and 200 µL of respective host bacterial culture (10^7^ cfu/ml) was mixed into 3 mL soft agar, poured on solid agar plate, and incubated up to 24 h at 37°C. Subsequently, SM buffer (4 mL) was gently poured on confluently lysed top agar. Plates were further incubated at 37°C for 4 hours with gentle shaking. Top agar and the remains of SM buffer were scraped and centrifuged at 5400 × g (4°C). The supernatant was filtered through 0.2 µm pore CA syringe membrane filters (Filtropur, Starsted). Prepared phage suspension was transferred to U-formed centrifuge tubes suitable for ultra-centrifugation (25 000 × g, 120 min, 4 °C) (Beckman Coulter, Optima^™^ MAX-XP). Pellets were resuspended in 200 µL of SM buffer and phage stock suspensions were stored at 4°C and −80°C.

Transmission electron microscopy was performed at National institute for biology, Ljubljana, Slovenia).

Host range of isolated phages was tested with the double-agar-layer assays using 12 *Bacteroides* strains belonging to four species (Data set S1).

### Lysogen formation assay

Each isolated phage was cultivated with its respective host strain. Plates with formed plaques in plaque assay were incubated in anaerobic chamber at 37°C for additional 72 hours, to allow the growth of potential lysogenic strains. From the plaques formed on double-layer agar, bacterial cultures were isolated with a sterile needle or small pipette tip and inoculated on sABB agar plates to obtain pure cultures. At least 12 strains were isolated per tested bacteriophage. Sensitivity of obtained strains for isolated phages was tested with DAL spot assay described above (Figure S4).

### Phage and bacterial genome sequencing

Phage lysate (200 µL) with app. 10^9^ pfu/ml was treated with DNase I (Sigma Aldrich) at the final concentration of 0.02 mg/ml and 0.05 mg/mL RNAse A (Sigma Aldrich) and incubated for 2h at 37°C, followed by 10 min heat inactivation at 90°C. Potential presence of host genomic residues was assayed with PCR using primers targeting 16S rRNA gene ^23^. Phage DNA was extracted with RTP^®^ DNA/RNA Virus Mini Kit following manufacturer’s instructions (INVITEK Molecular).

Bacteroides DNA was extracted (QIAamp DNA Mini Kit, Qiagen).

For phage and bacterial genomes paired-end libraries were generated using the Nextera XT Library preparation kit (IIlumina) and sequenced on MiSeq (Ilumina) with 600-cycle MiSeq ReagentKit v3.

The quality of the raw sequencing reads was examined by FastQC tool Version 0.11.9 (Babraham Bioinformatics) ^25^. Quality trimming was done by Trimmomatic Version 0.39 (USADELLAB.org) ^26^ and overlapping paired-end reads were merged by using FLASH software, version 1.2.11 (CBB) ^27^. Assembly was performed by SPAdes Assembler, version 3.14.0 28 and the assemblies were examined using Quast version 4.0 ^29^. Genomes were then annotated with Prokka 1.14.5 ^30^.

### Bacteriophage genome annotation

Protein sequences of open reading frames (ORFs), determined with Prokka 1.14.5 ^30^, were blasted (blastp, NCBI, 2019) against non-redundant protein sequences (nr) database. Conserved protein domains of ORF were predicted with Conserved Domain Search (CDD, NCBI) and Pfam ^31^. Additionally, remote homologues were also detected using PHYRE2 – Protein Homology/analogY Recognition Engine V 2.0 ^32^. Presence of signal peptides was analyzed with SignalP-5.0 Server ^33^. Remote homologs of phage head-neck-tail module proteins were additionally analyzed on VIRFAM sever ^34^. Predicted DGR regions were analyzed with myDGR, a server for identification and characterization of diversity-generating retroelements ^35^.

### Phage classification and phylogenetic analysis

Phage life style and classification was computationally analyzed using PHACTS program (http://www.phantome.org/PHACTS/index.htm) ^36^.

vConTACT2 ^37^ was used for taxonomic classification using the ViralRefSeq-prokaryotes-v94 database. To determine phage DNA packaging and replication strategy, a phylogenetic analysis of amino acid sequences of TerL – terminase large subunit was made. Sequences of TerL were downloaded from NCBI and Pfam databases and aligned using the ClustalW ^38^ program. Phylogenetic tree was then generated with the SeaView Version 5.0.2 ^39^ integrated phyML using the maximum likelihood approach and GTR nucleotide substitution model. The resulting dendrogram was then visualized with FigTree v1.4.4 (http://tree.bio.ed.ac.uk/software/figtree/).

### Identification of shared homologous proteins and prophage regions

Based on closest BLASTp hits of determined ORFs, closest relatives were manually predicted and their bacterial host genomes were examined for prophage presence. Ranges of prophage regions were determined based on the G+C content, predicted functional annotations of neighboring genes, presence of integrase and other phage specific genes or identification of repeats sites (attL and attR). Sequences of predicted prophage regions were extracted from host genomes using Artemis software version 1.8 ^40^, annotated with Prokka 1.14.5 ^30^ and applied in comparison using Easyfig ^41^. Protein sequences of ORFs of identified prophages were analyzed for conserved protein domains like described above. Gene synteny in different phage functional gene groups was analyzed.

### SNP analysis of potential phage target genes

Reads of original phage host (*B. uniformis* MB18-80) and two derivative strains isolated in lysogeny experiment (MB18-80-K and MB18-80-PH) were mapped to original MB18-80 assembly using BBTools ^42^. Sorted BAM files were used for calling SNPs sites using the SAMtools verison 0.1.19 ^43^. Mapped reads and SNP sites were also analyzed relative to MB18-80 genome using Artemis software version 1.8 ^40^.

### Tandem repeats analysis with direct sequencing

Tandem repeats were located and analyzed with Tandem Repeats Finder ^44^. Primers (primer F2, 5’-CCTCGGTAATGCTTTCTACG-3’; primer R2, 5’-AGGTAGCCGTAAATGTATCG-3’) were constructed using SnapGene software (GSL Biotech LLC, 2004) and were used in a direct Sanger sequencing reaction (40 cycles; using a gDNA as a template and BigDye Terminator v3.1 Cycle Sequencing Kit) to examine if the repeats represent phage genome termini of linear dsDNA phage. Sequencing was performed on 3500 Series Genetic Analyzer (ThermoFisher Scientific) and analyzed with Artemis software version 1.8 ^40^

### Metagenomic analysis

Paired-end sequencing reads in fastq format of metagenomics studies under the BioProject accession numbers PRJNA491626, PRJNA268964 and PRJNA278393 were downloaded from The European Nucleotide Archive (ENA) (https://www.ebi.ac.uk/ena). Adaptor removal and quality trimming was conducted by Trimmomatic Version 0.39 (USADELLAB.org) ^26^. Processed metagenomics reads were mapped to genome assembly of isolated phage using BBTools ^42^.

### Data availability

The assembled genomes were submitted to the NCBI (https://www.ncbi.nlm.nih.gov/) under the Bioproject accession numbers PRJNA636979 (bacterial genomes) and PRJNA638235 (phage genomes).

## Results

### Isolation and phenotypic characterization of phages specific for Bacteroides uniformis

In 8 out of 12 *Bacteroides* strains belonging to four species the lysis like zones were observed. Subsequently, plaques were successfully propagated from two *B. uniformis* strains (Data set S1). Circular plaques were formed with diameter ranging from 0.1 to 3 mm (Figure 1, B). Four seemingly different bacteriophages were isolated (F1-F4). Phages were stable if stored at 4°C or - 80°C, at high concentration (10^11^ pfu/ml). Subsequent analysis showed that phages F3 and F4 were genetically identical and thus for further experiments only phages F1, F2 and F4 were used. Host range was tested on all *Bacteroides* strains included in this study (Data set S1). In addition to the initially identified *B. uniformis* host strains, lysis like forms (Figure 1, C) were observed with additional representatives of *B. vulgatus, B. uniformis, and B. ovatus*, although we were not able to further propagate the phages.

**Figure 1.**
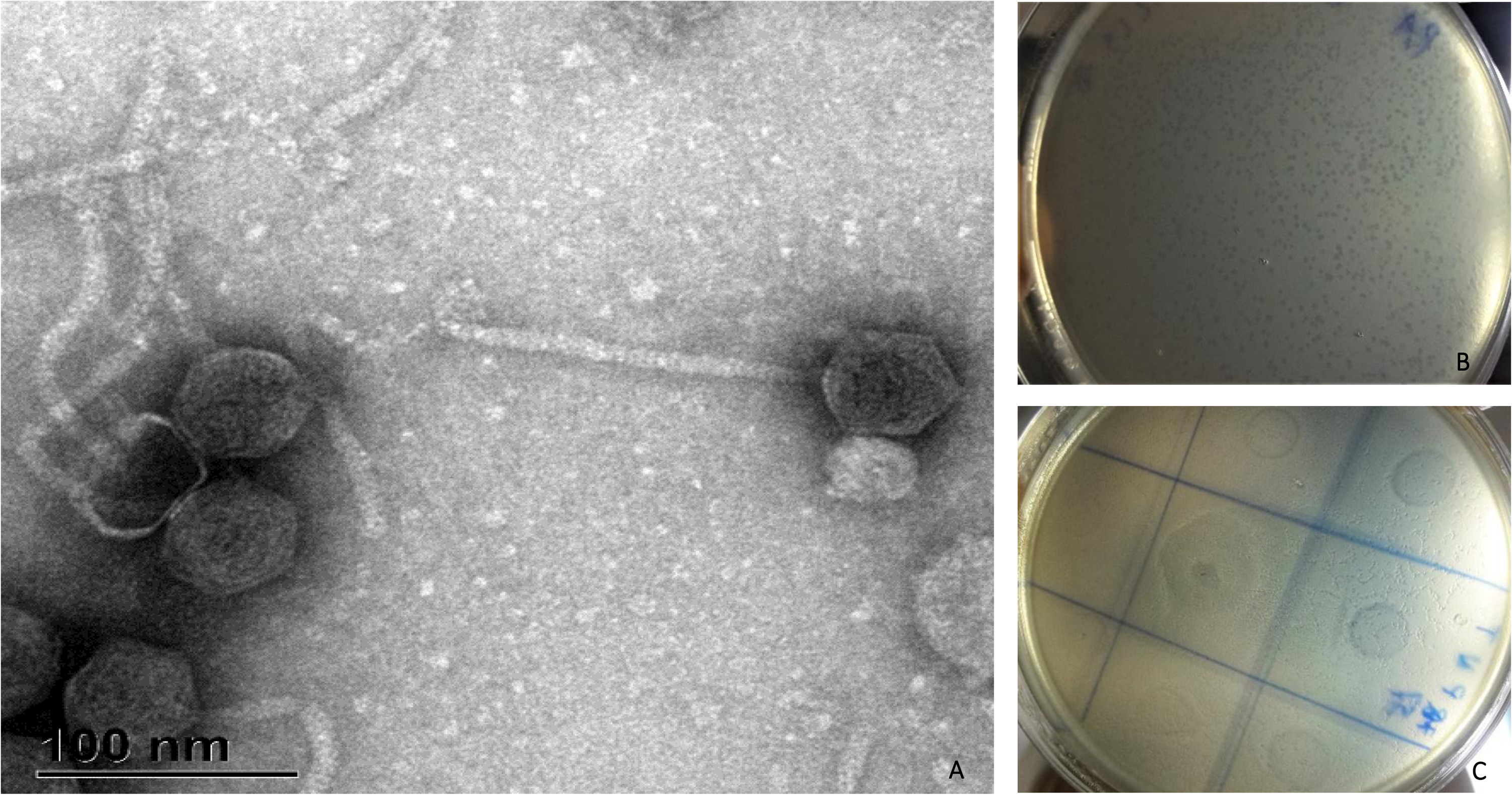
Lytic Bacuni phages exhibit *Siphoviridae* morphology. (a) Photograph of Bacuni virion obtained by transmission electronic microcopy (scale bar is 100 nm). (b) Plaque morphology of Bacuni phage F1 formed on *B. uniformis* MB18-33 host lawn after 24 hours incubation in sABB agar overlay. (c) Lysis like zones formed on sABB agar overlay after 24 incubation with host strain *Bacteroides vulgatus* MB18-32 in double layer agar overlay (spot assay with enrichment sample).

Attempts to isolate potential lysogenic *Bacteroides* strains from the formed plaques were not successful. Only 10 out of 35 inoculated plates resulted in bacterial growth. These strains were further tested for susceptibility to infection with obtained phages. Experiment was performed three times and no lysogens were detected (discussed in detail below).

Transmission electron microscopy (TEM) analysis showed morphology typical of the *Siphoviridae* family of the *Caudovirales* with icosahedral heads of about 50 nm in diameter and approximate tail size of 150×8 nm (Figure 1, A).

### Novel B. uniformis phages show high degree of similarity to each other and belong to a new genus

The assembled genome lengths of phages F1, F2 and F4 were from 40421 to 40653 bp (Table 1). G+C content of phage genome content was 51.8 mol % (F1), which is considerably higher than its host genome G+C content (46.3 mol %), obtained from WGS analysis, which is also consistent with *Bacteroides uniformis* reference stain G+C content ^44^.

**Table 1.** Comparison of general characteristics of isolated phages belonging to a newly defined genus Bacuni.

| Phage | F1 | F2 | F3 and F4 |
| --- | --- | --- | --- |
| Bacterial host | *Bacteroides uniformis*<br>MB18-33 | *Bacteroides uniformis*<br>MB18-33 | *Bacteroides uniformis*<br>MB18-80 |
| No. of predicted ORFs | 51 | 51 | 50 |
| Assembled genome length (bp) | 40421 | 40653 | 40640 |
| G+C content (%) | 51.8 | 51.7 | 51.7 |
| Genetic differences (compared to F1) | Reference | 16 SNPs in the DGR and 19 aa insertion in RT gene | 24 SNPs, 18 in DGR<br>RT identical to RT F1 |

(RT – reverse transcriptase, VR – variable repeat, aa – amino acids, SNP – single nucleotide polymorphism, ORF – open reading frame; DGR: Diversity-generating retroelements)

All four isolated phages were similar one to another (99.83 % similarity) (Table 1). Genomes of phage F1 and F4 differ only in 24 SNP sites, of which 18 are condensed in variable repeat region 1 (VR 1) of DGR and the phages infect different hosts. Phage F2 shares the same host with phage F1 but deviates from F1 in an insertion of 19 aa in putative reverse transcriptase gene of the DGR and in 16 SNPs in variable repeat regions (VR) of the DGR.

The isolated phages could not be assigned to any of the known prokaryotic viral clusters using a gene sharing network approach vConTACT2 ^37^, implying that so far no similar bacteriophages have been reported (Data set S2 (A) and Figure S1 (B)). Based on no resemblance with phage genera described to date, phages F1, F2 and F4 were classified as a new genus, and for the purpose of this paper provisionally named Bacuni.

TEM based classification of Bacuni phages into *Siphoviridae* family was additionally confirmed *in silico* using Virfam server ^34^, which identifies proteins of the phage head-neck-tail module and assigns phages to the most closely related cluster of phages within the ACLAME ^45^ database (Figure S2).

### Genome organization of novel B. uniformis phages

Using automated annotation, 51 open reading frames (ORFs) were predicted in Bacuni genomes. Further functional annotation lead to a prediction of potential functions of 34 genes, which could be divided in five common phage functional groups (Figure 2 and Data set S3).

**Figure 2.**
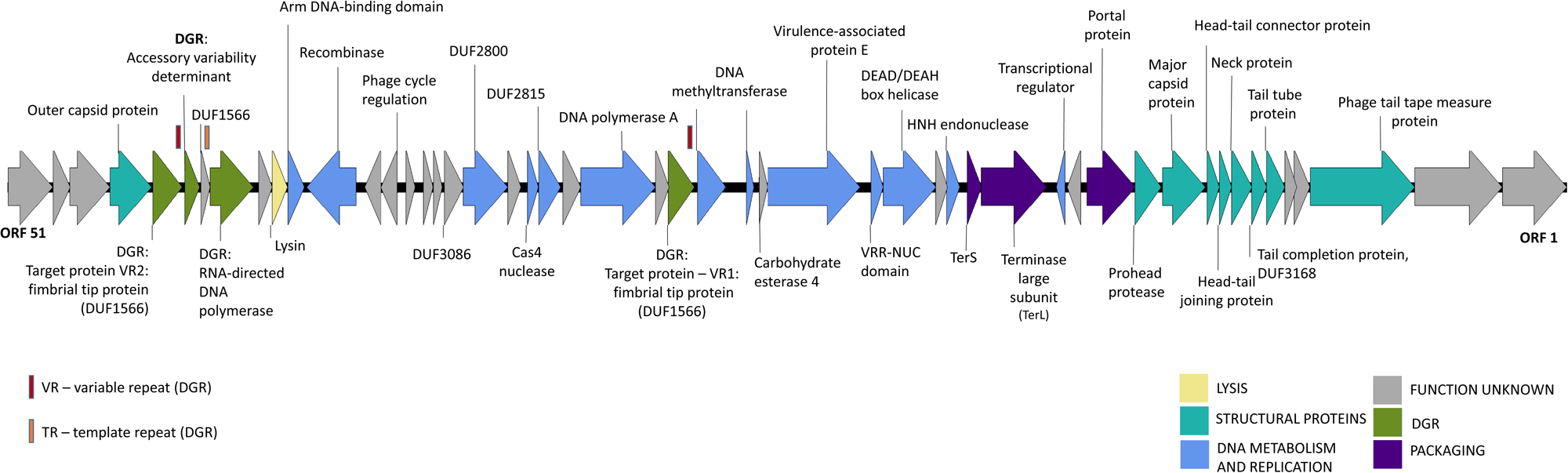
Linear genome map of Bacuni phage F1. Colors of open reading frames correspond to the general predicted functions (see color legend for details). Genes with no functional annotations (hypothetical proteins) are not labeled. Locations of template sequence (TR) and variable repeats of diversity-generating retroelement (DGR) are marked with orange and red rectangles above associated proteins.

Tandem nucleotide repeats were identified in Bacuni phage ORF for putative phage tail tape measure protein and direct sequencing was conducted to examine whether repeats in phage genome are terminal, which was not the case. Phylogenetic analysis of large terminase subunit genes (TerL) (Figure S3) indicated that Bacuni phages use rolling circle-concatemer genome replication due to clustering into the group of phages with cohesive ends and 3′-single-strand extensions.

Nine putative structural proteins were identified, including the major capsid protein, prohead protease, and a large phage tail tape measure protein with observed tandem repeats typical for these proteins ^46^ and four transmembrane helices. Large and small subunit of the terminase and portal protein, which together form a packaging function group, were found located in the close proximity of the structural genes. Bacterial cell wall hydrolytic enzyme, a predicted acetylmuramoyl-L-alanine amidase, was identified as a putative lysin.

Based on conserved domain search, twelve identified phage genes are putatively involved in DNA metabolism and replication. Additionally, two genes primarily identified as Domains of unknown function (DUF2800 and DUF2815) were recently assigned new putative roles by bioinformatic approach ^47^. They are likely to be involved in regulation of phage DNA metabolism. DUF2815 hypothetically functions as single-stranded DNA and DUF2800 as a cis-regulatory elements or small RNA in phages ^47^.

Finally, four functionally annotated genes belong to diversity-generating retroelement (DGR).

### DGR variability and host tropism

Diversity-generating retroelements are recently described genetic elements that use reverse transcription from a donor template repeat (TR) to a recipient variable repeat (VR) in defined target gene. This generates vast numbers of sequence variants (substitutions) in specific target genes ^10^.

VR sequences of Bacuni phages are located on genes whose products exhibit DUF1566 and/or Fib_succ_major motifs. The Legionella DGR exemplifies the closest studied DGR ^19,48^. DGRs found in Bacuni phages belong to a group operating on targets exhibiting a C-type lectin fold ^19^. This classification and the presence of DGR elements in Bacuni phages were also confirmed with MyDGR, a server for identification and characterization of diversity-generating retroelements ^35^.

Bacuni phages have two target genes putatively diversified by DGR. First target gene (with detected VR 2) is located on a distant part of the phage F1 genome (6947-7054 bp) while the second target gene with VR 1 (20281-20388 bp) is found in the immediate neighborhood of the core DGR components including reverse transcriptase (RT) (18141-19544 bp), Avd-accessory protein (19814-20197 bp), and the TR containing gene (19577-19684 bp). The variable repeat gene region, which is diversified, lies at the 3’ end of target genes and codes for the last 35 amino acids. Both variable repeats were found at 3’ end of the target gene with DUF1566 domain, where also almost all genetic differences between Bacuni phages are located (Table 1, Figure 3). Bacuni phages F1 and F4 differ in 13 amino acids in this region. Target genes in Bacuni phage genomes were found in ORFs that include motifs for cellular adhesion and represent a putative fimbrial tip protein. Identified target genes exhibit high similarity with 60 % or higher coverage (Phyre2) to crystal structure of a fimbrial tip protein (bacova_04982) from *Bacteroides ovatus* atcc 8483 ^49,50^ that was also identified as a DGR target in metagenomes of human stool samples ^49^

**Figure 3.**
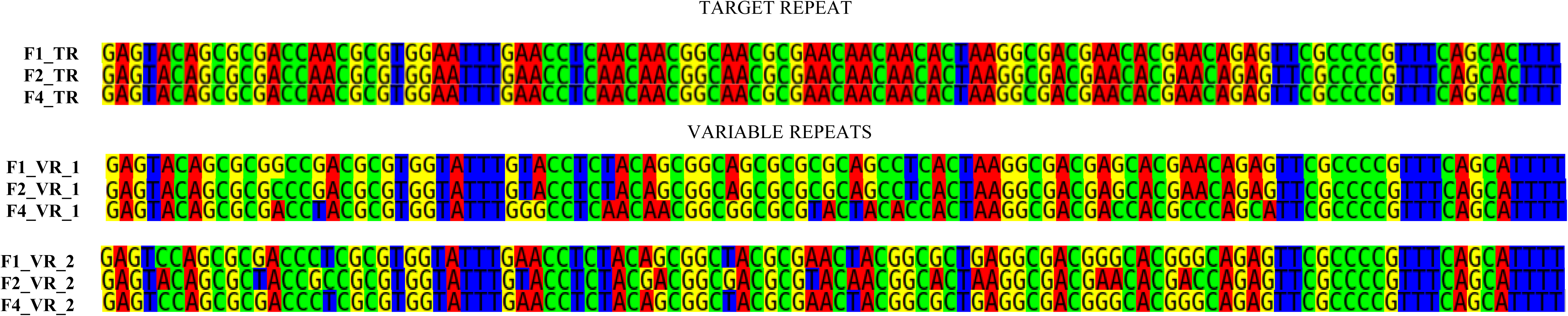
Alignment of the TRs and VRs from isolated Bacuni phages. Each nucleotide base is color-coded for visualization of mismatches in the variable repeat. VR2, located in the close proximity of reverse transcriptase represent the region with highest condensation of SNP sites, which most likely influence Bacuni phage host range.

The observed TR-VR substitutions can be seen in Figure 3 and are, as expected, mutations in adenines. They are most probably the results of induced substitutions mediated by RT (Figure 3).

Despite high genetic similarly, isolated Bacuni phages exhibit different host range (Table 1). Since the vast majority of genetic differences was concentrated in VR regions of DGR target genes located in putative fimbrial tip proteins, we propose that DGRs influence host range of Bacuni phages.

### Bacuniphage similarities with other phages and prophages of various anaerobic bacteria

As described above, searches against the NCBI non-redundant database and the Reference Viral Database ^51^ showed no similarities of Bacuni phages to any known phages at the nucleotide level. BLASTp search, however, revealed some homology to prophage-related gene products encoded in the genomes within the order *Bacteroidales* (Table 2; Figure 4).

**Table 2.** Comparison of selected genome characteristics between Bacuni phages and putative partially homologues prophage genomes

| Host strain | Source | Collection date and location | Region length (bp) | No. of Bacuni homologous proteins / No. of ORFs | Coverage (%) - nt identity (%) * | Genome location and biosample accession |
| --- | --- | --- | --- | --- | --- | --- |
| *Bacteroides acidifaciens* NM70_E10 | *Mus musculus*, colon and cecum | 2016, Toronto, Canada | 44986 | 28 / 48 | 45% - 71.06% | Node 8 (64227 ... 109212)<br>SAMN10878312 |
| *Prevotella* sp. P3-122 | *Sus scrofa domestica*, feces | 2014, Slovenia: pig farm Ihan | 34280 | Contig 46: 21 / 35,<br>Contig 76: 5 / 15 | 46% - 72.82% | Contig 46 (44340 ... 78619)<br>Contig 76 (1 ... 12007)<br>SAMN07431220 |
| *Prevotella* sp. OH937_COT-195 | *Canis lupus*, dog mouth | 2012, Leicestershire, UK | 38640 | 17 / 47 | 28% - 71.82% | Scaffold20 (3163 ... 37390)<br>SAMN10478691 |
| *Porphyromonas gingivicanis* COT-022 OH1391 | *Canis lupus*, dog mouth | 2012, Leicestershire, UK | 35922 | 11 / 39 | 23% - 70.63% | Contig 6 (16379 ... 52300)<br>SAMN03004338 |
| *Porphyromonas cangingivalis* JCM 15983 | n.a. | 2014, The University of Tokyo | 33481 | 17 / 46 | 33% - 67.78% | Node 1 (3106 ... 36586)<br>SAMD00003336 |
| *Prevotella timonensis* UMB0818 | *Homo sapiens*, catheter | 2015, USA: Maywood, IL | 37867 | 16 / 47 | 4% - 69.19% | Node 1 (56177 ... 94272)<br>SAMN07511428 |

\*Genome coverage ~ Percent of nucleotide identity (discontiguous megablast) compared to F1

**Figure 4.**
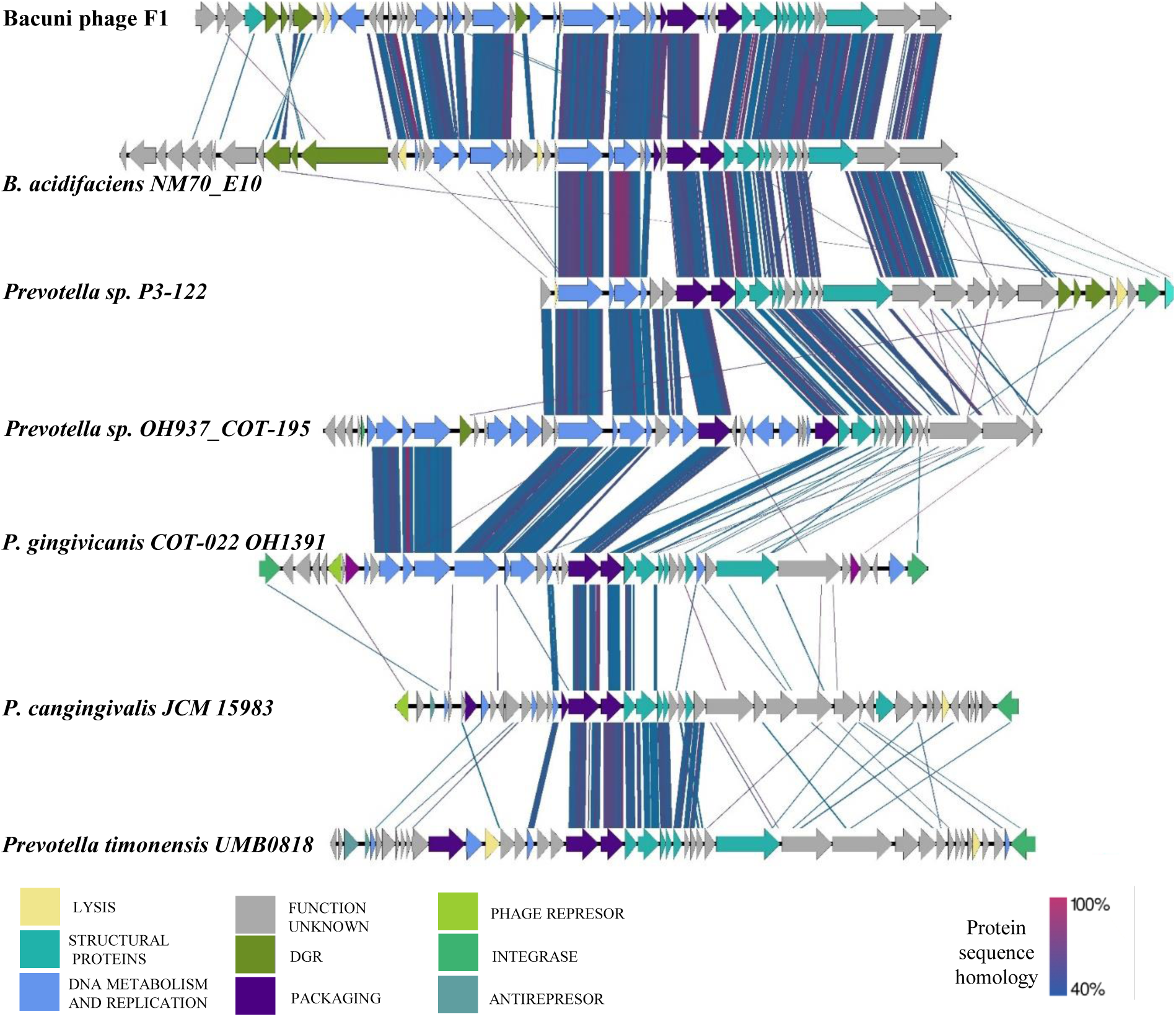
Comparison of genome organization and genomic synteny of Bacuni phages to putative prophage genomes in various bacterial hosts from *Bacteroidales*. BLASTp sequence homology (40 % similarity and higer) between Bacuni phage F1 and related prophage regions identified in genomes of *B. acidifaciens, Prevotella sp*., *P. gingivicanis* and *P. cangingivalis* (see Table 2 for more information) is indicated with a color link. Colors of putative proteins correspond with the general predicted functions (see color legend).

Six putative prophage regions were identified in assembled bacterial genomes with reliable homologies (Table 2; Figure 4).

Some of the identified prophage regions were found on contig borders and some assemblies were highly fractioned, thus some parts of prophage genomes could have been left out. The putative functions of retained prophage ORFs were assigned based on conserved protein domains found (Data set S4). The identified putative prophage regions have not been described before.

The highest homology (up to 85% amino acid similarity) to proteins of Bacuni phages was observed in putative prophage regions of *B. acidifaciens* NM70_E10 and *Prevotella* sp. P3-122 (Figure 4). They share significant protein homology between two thirds of annotated proteins of various functional clusters including the DGR region and its target region overlapping DUF1566 domain. However, no homologies were found in its putative lysin and recombinase genes.

Protein level homologies found in remaining identified putative prophage regions of *Prevotella* sp. OH937_COT-195, *Porphyromonas gingivicanis* COT-022 OH1391, *P. cangingivalis* JCM 15983 and *Prevotella timonensis* UMB0818 were mostly present in structural and packaging functional gene groups (Figure 4).

The prevalence of predicted prophage regions identified in initial screening (Table 2) was further examined in Genebank nr-database. Minor nucleotide level similarities of the predicted prophage regions were found, with a few exceptions. Nucleotide homology (92%) on 30% of putative *B. acidifaciens* NM70_E10 prophage region length was found in genomes of *B. ovatus* 3725 D1 (CP041395.1), *Bacteroides xylanisolvens* strain H207 (CP041230.1), and in unidentified phage clone 1013 (JQ680349.1).

Whole sequence of predicted *P. cangingivalis* JCM 15983 prophage was also found in the genome of *P. cangingivalis* ATCC 700135 isolated in Finland and in *P. cangingivalis* NCTC12856 collected in 1986 and isolated from fecal sample of *Homo sapiens*.

### Identification of Bacuni phages in Human gut virome database and in associated metagenomes

Genome of Bacuni phage F1 was blasted (blastn) against Human gut virome database (GVD), a novel database composed of 13,203 unique viral populations obtained from gut metagenomes of 572 individuals from different geographical locations ^52^. Matches (roughly 80% nucleotide similarity over more than 80% of the Bacuni phages) were found in contigs originating from two studies ^53,54^. Data was further tracked to authentic metagenomics data sets that include metagenomes from Western urban societies and traditional communities ^53,54^. Search for reads mapping to Bacuni phage genome revealed that Bacuni phages were underrepresented in Western data sets analyzed, but present in data sets of fecal viromes of Cameroonians with gastroenteritis (Data set S5). Up to 6066 reads from metavirome of a Cameroonian ^54^ were found to align to Bacuni phage, majority originating from the Kumba region (Data set S5). Further analysis showed that those reads cover 31 of the 40 kb Bacuni phage F1 genome.

### Changes of host susceptibility pattern after exposure to Bacuni phage

Assay for detection of lysogenic bacteriophage in *B. uniformis* host strains was conducted (Data set S1, Figure S4). Three attempts to isolate potential lysogenic host strains from the formed plaques resulted each in roughly 10 viable derivatives of *B. uniformis* MB18-80 and *B. uniformis* MB18-33. These derivative strains were retested with all three Bacuni phages. Spot assay showed mixed results: some derivatives were indeed not lysed by any of the phages (representing possible lysogens), while some were resistant to challenging phages but lysed by phages that initially did not lyse the original strain. Thus, this was not a simple lysogenization.

Two derivatives of *B. uniformis* MB18-80, host of phage F4, were further selected for WGS: MB18-80-K, a potential lysogen, that was resistant to infection with all tested phages, and second derivative MB18-80-PH that became susceptible to infection with phages F1 and F2, but resistant to F4 (Data set S1, Figure S4).

Genome analysis of *B. uniformis* MB18-80-K and *B. uniformis* MB18-80-PH disproved assumptions of lysogenic lifestyle since no parts of Bacuni phage genome were detected in genome of sequenced derivative strains. These results were in agreement with the predicted lytic life style of isolated phages with Phage classification tool set (PHACTS)^36^.

Comparison of the obtained *B. uniformis* derivative genomes to original host strain indicated SNPs in several biologically relevant genes (Table 3, Data set S6). Genome of immune derivative *B. uniformis* MB18-80-K exhibits SNPs in genes coding for putative restriction enzymes involved in defense mechanism against invading viruses and in outer membrane transporter complexes most likely involved in import of large degradation products of proteins or carbohydrates (Table 3, Data set S6). *B. uniformis* MB18-80-PH, in which phage tropism switching was observed, exhibited SNPs in partially overlapping set of genes coding for restriction enzymes, putative porins, peptidoglycan binding proteins, and putative peptidoglycan hydrolase (Table 3, Data set S6).

**Table 3.**
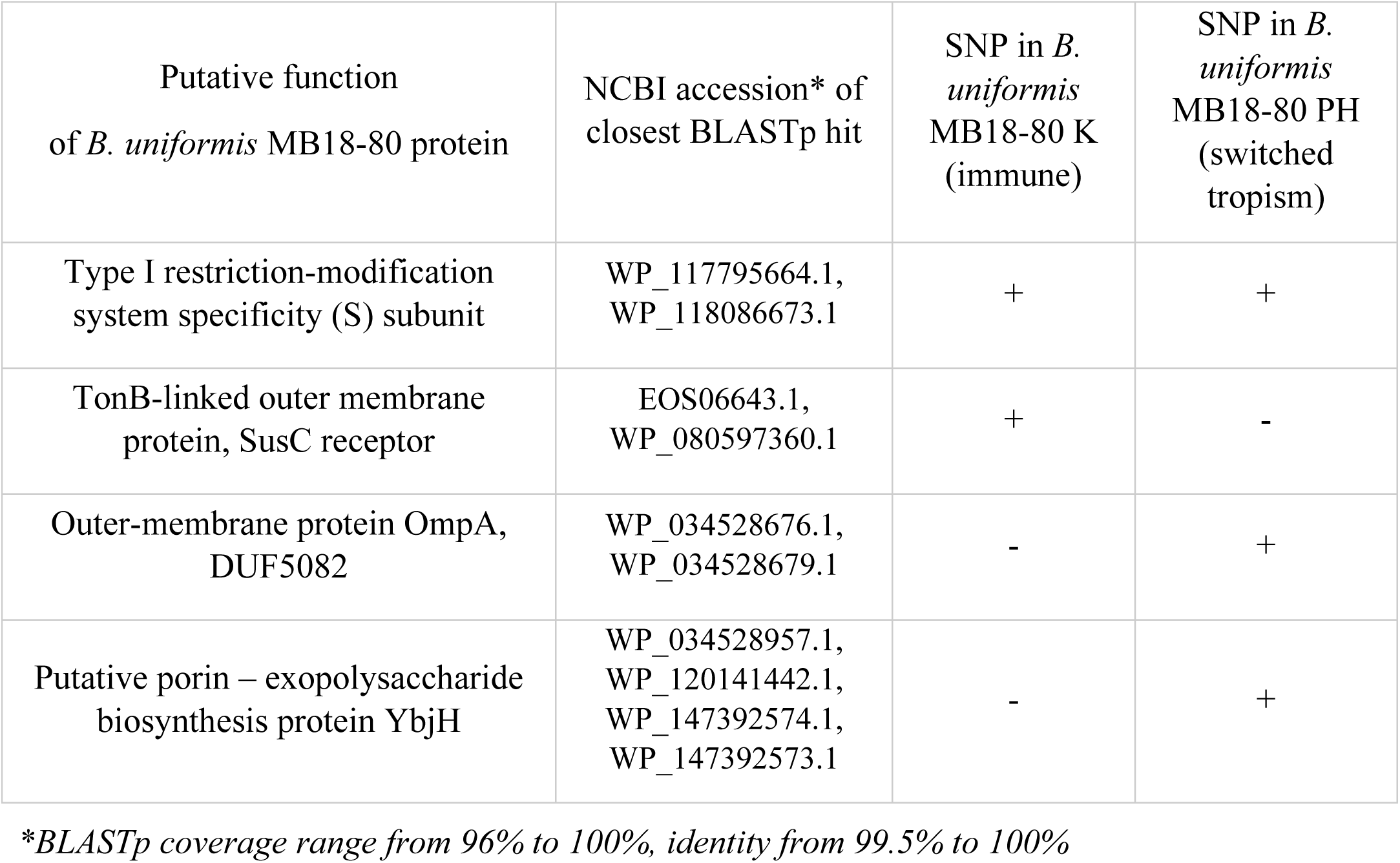
Genetic differences in biologically relevant genes of Bacuni phage F4 host MB18-80 and its derivatives that are immune to infection with Bacuni phages or indicate tropism switching pattern.

| Putative function<br>of *B. uniformis* MB18-80 protein | NCBI accession* of<br>closest BLASTp hit | SNP in *B.*<br>*uniformis*<br>MB18-80 K<br>(immune) | SNP in *B.*<br>*uniformis*<br>MB18-80 PH<br>(switched<br>tropism) |
| --- | --- | --- | --- |
| Type I restriction-modification<br>system specificity (S) subunit | WP_117795664.1,<br>WP_118086673.1 | + | + |
| TonB-linked outer membrane<br>protein, SusC receptor | EOS06643.1,<br>WP_080597360.1 | + | - |
| Outer-membrane protein OmpA,<br>DUF5082 | WP_034528676.1,<br>WP_034528679.1 | - | + |
| Putative porin – exopolysaccharide<br>biosynthesis protein YbjH | WP_034528957.1,<br>WP_120141442.1,<br>WP_147392574.1,<br>WP_147392573.1 | - | + |

\*BLASTp coverage range from 96% to 100%, identity from 99.5% to 100%

## Discussion

*Bacteroides* is one of the most prominent bacterial genera of the human gut microbiome and is known as dietary fiber fermenter that produces short chain fatty acids important for host health ^55,56^. As such it is commonly found in globally conserved core gut microbiota ^57–60^.

In this study, we describe isolation and characterization of human gut associated phages infecting *B. uniformis*. As they were essentially not similar to any of the hitherto described phages based on their encoded proteins, we were not able to classify them using VconTACT2. Thus, they may be the first isolated representatives of a new phage genus, provisionally named here a “Bacuni phage”.

Three isolated phages infected distinct *B. uniformis* strains. The tropism of Bacuni phages appears to be dependent on interplay of DGR mediated sequence variations of phage fimbrial tip proteins and mutations in host genes coding for outer-membrane proteins. Different host range between genetically very similar Bacuni phages can be explained with SNPs sites condensed in variable repeat of DGR region, located in a putative fimbrial tip protein, a gene presumably involved in cell adhesion and possibly acting as a cell receptor in Bacuni phages. The SNPs are at the C-terminus of two target proteins, at variable repeats that each consist of 35 amino acids. Bacuni phages F1 and F4 differ in 13 amino acids at the variable repeat 2 coded fimbrial protein tip end and infect different hosts, while Bacuniphages F1 and F2, that infect the same host, differ in variable repeat 1 far removed from the DGR region. Given that there are only 6 more SNPs observed between F1 and F4 outside of DGR, one may conclude that variable repeat 2, located in close proximity of reverse transcriptase, presumably plays a decisive role in Bacuni phage tropism in our experimental setting. These findings correlate with study where metagenomics data set from Human microbiome project (HMP) was screened for DGRs ^50^. There, the identified variable regions were also localized in a DUF1566 domain coding genes and the target protein showed high protein homology to a pilin tip from *Bacteroides ovatus* ^11,49^.

It appears that DGR contributes to increased adaptability of temperate and lytic phages in such complex communities as the human gut, where multiple species of the same genus and several strains of the same species may coexist. This evolutionary advantage may (indirectly) affect microbial diversity and influence health of the associated mammalian host.

To the best of our knowledge, Bacuni phages represent the first DGR-containing lytic phages ^11^. Based on protein homologies to here described six putative prophages (Figure 4, Table 2) and their paucity in virome studies, it is plausible that Bacuni phages originate from temperate phages.

Viral databases do not contain many genomes of phages infecting dominant gut bacteria and we were initially not able to locate a metagenome/virome that contained sequence reads mapping to Bacuni phages. However, recently published GVD database improves viral detection rates over NCBI viral RefSeq by nearly 60-fold” ^52^. Almost complete Bacuni phage genome was found in GVD originating from intestinal viromes of Cameroonians ^54^. Weak signal of reads in metagenomes of traditional Peruvian communities and urban Italian gut metagenomes ^53^ may indicate that these phages are present at various geographic locations but not abundant enough to be detected with common metagenomics sequencing technologies that are generally not yet optimized to detect bacterial viruses.

Our study sheds light on feasibility of isolation of lytic phages infecting abundant gut bacteria. Lytic phages are suitable for use in phage therapy ^61–64^. In vivo studies in mice using commercial phage cocktails showed that phages triggered a cascade reaction that influenced bacterial diversity and composition ^65^. Additional further research may provide phages targeting less beneficial bacteria in the intestine with potential therapeutic role on human gut microbiota.

In summary, phages described in this study represent a new genus, are the first example of phages using *B. uniformis* as a host, are one of the rare lytic phages isolated from the gut ecosystem and are the first lytic phages with DGR sequences. Single nucleotide variation in phage DGRs and in the relevant host proteins are described in the context of host specificity pattern changes.

## Acknowledgements

Authors would like to thank Magda Tusek Znidaric for performing TEM.

## Supplemental Material

Data set S1

List of *Bacteroides* strains, isolated from stool sample and associated phage screening and host range experiments.

Data set S2 (A) and Figure S1 (B)

Taxonomic anaylsis conduced with vConTACT2 ^37^ shows that isolated phages could not be assigned to any of the known prokaryotic viral clusters. Supplemental file S2 contains the Cytoscape network file (B) and the data set (A) with viral clusters made by vConTACT2. In the file the phage F1 is named 3P11 and the phage F4 8POS.

Figure S2

Classification of the Bacuni phage F1 with respect to other related phages in Aclame (Bacuni phage F1 in text box with red border and white background). According to Virfam server generated protein identification of the phage head-neck-tail module, Bacuni F1 clusters into *Siphoviridae* of the neck type 1, cluster 3 within the phages in the database ACLAME. The conserved genome organization observed among the phages of the ACLAME database was used to define allowed inter-gene distance intervals ^34^. Each cluster with associated number represents a different neck type.

Data set S3

Putative functions of identified ORFs of Bacuni phages and their closest BLAST hits. Functional annotations for each Bacuni F1 predicted ORF. Function were determined by comparisons to the conserved domain database, Pfam, Phyre2, Virfam and MyDGR. For each ORF best BLASTp hit accession with the corresponding e-vaule, query coverage and percent identity is listed. Predicted signal peptides and transmembrane domains are included.

Figure S3

Phylogenetic analysis of terminase large subunit (TerL) generated with phyML using the maximum likelihood approach and GTR nucleotide substitution model. Bacuni phage TerL clusters into the group of phages with cohesive ends and 3′-single-strand extensions

Data set S4

The putative functions of ORFs encoded in 6 identified prophages that share homologous proteins with Bacuni phages. Functions were determined by comparisons to the conserved domain database, Pfam, Phyre2, Virfam and MyDGR. For each ORF best BLASTp hit accession with the corresponding e-vaule is listed

Data set S5

Number of aligned reads mapping to Bacuni phage F1 in human gut derived metagenomics data sets. The Bacuni phage F1 genome was used as a reference to align reads from whole-community metagenomes using BBtools.

Figure S4

Schematic methodologic overview of lysogenic assay conducted to explore Bacuni phage host range.

Data set S6

Genetic differences in biologically relevant genes of Bacuni phage F4 host MB18-80 and its derivatives that are immune to infection with Bacuni phages or switched host tropism. Genome location of genes with SNPs and their closest blastp hit accession numbers are provided.

## References

1. Sausset R, Petit MA, Gaboriau-Routhiau V, De Paepe M. New insights into intestinal phages. Mucosal Immunol. 2020;13(2):205–215. doi:10.1038/s41385-019-0250-5

2. Sordi L De, Lourenço M, Debarbieux L. “I will survive “: A tale of bacteriophage-bacteria coevolution in the gut. Gut Microbes. 2019;10(1):92–99. doi:10.1080/19490976.2018.1474322

3. Aggarwala V, Liang G, Bushman FD. Viral communities of the human gut?: metagenomic analysis of composition and dynamics. 2017:1–10. doi:10.1186/s13100-017-0095-y

4. Minot S, Sinha R, Chen J, et al. The human gut virome?: Inter-individual variation and dynamic response to diet The human gut virome?: Inter-individual variation and dynamic response to diet. 2011:1616–1625. doi:10.1101/gr.122705.111

5. Minot S, Bryson A. Rapid evolution of the human gut virome. Proc …. 2013;110(30):12450–12455. doi:10.1073/pnas.1300833110//

6. Dutilh BE, Cassman N, McNair K, et al. A highly abundant bacteriophage discovered in the unknown sequences of human faecal metagenomes. Nat Commun. 2014;5:1–11. doi:10.1038/ncomms5498

7. Guerin E, Shkoporov A, Stockdale SR, et al. Biology and Taxonomy of crAss-like Bacteriophages, the Most Abundant Virus in the Human Gut. Cell Host Microbe. 2018;24(5):653-664.e6. doi:10.1016/j.chom.2018.10.002

8. Shkoporov AN, Khokhlova E V, Fitzgerald CB, et al. ΦCrAss001 represents the most abundant bacteriophage family in the human gut and infects Bacteroides intestinalis. Nat Commun. 2018;9(1):4781. doi:10.1038/s41467-018-07225-7

9. Andrew J. Hryckowian, Bryan D. Merrill, Nathan T. Porter W Van, Treuren, Eric J. Nelson, Rebecca A. Garlena, Daniel A. Russell EC, Martens JLS. Bacteroides thetaiotaomicron-infecting bacteriophage isolates inform sequence-based host range predictions. preprint. 2020. doi:10.32388/nqwmca

10. Cornuault JK, Petit M, Mariadassou M, et al. Phages infecting Faecalibacterium prausnitzii belong to novel viral genera that help to decipher intestinal viromes. 2018:1–14.

11. Benler S, Cobián-Güemes AG, McNair K, et al. A diversity-generating retroelement encoded by a globally ubiquitous Bacteroides phage 06 Biological Sciences 0605 Microbiology. Microbiome. 2018;6(1):1–10. doi:10.1186/s40168-018-0573-6

12. Devoto AE, Santini JM, Olm MR, et al. Megaphages infect Prevotella and variants are widespread in gut microbiomes. Nat Microbiol. 2019;4(4):693–700. doi:10.1038/s41564-018-0338-9

13. Cornuault JK, Moncaut E, Loux V, et al. The enemy from within: a prophage of *Roseburia intestinalis* systematically turns lytic in the mouse gut, driving bacterial adaptation by CRISPR spacer acquisition. bioRxiv. January 2019:575076. doi:10.1101/575076

14. Hawkins SA, Layton AC, Ripp S, Williams D, Sayler GS. Genome sequence of the Bacteroides fragilis phage ATCC 51477-B1. Virol J. 2008;5(Figure 1):1–5. doi:10.1186/1743-422X-5-97

15. Ogilvie LA, Caplin J, Dedi C, et al. Comparative (meta)genomic analysis and ecological profiling of human gut-specific bacteriophage φB124-14. PLoS One. 2012;7(4):1–17. doi:10.1371/journal.pone.0035053

16. Gilbert RA, Kelly WJ, Altermann E, et al. Toward understanding phage: Host interactions in the rumen; complete genome sequences of lytic phages infecting rumen bacteria. Front Microbiol. 2017;8(DEC):1–17. doi:10.3389/fmicb.2017.02340

17. Pacífico C, Hilbert M, Sofka D, et al. Natural occurrence of Escherichia coli-infecting bacteriophages in clinical samples. Front Microbiol. 2019;10(OCT):1–18. doi:10.3389/fmicb.2019.02484

18. Sears CL. A dynamic partnership: Celebrating our gut flora. Anaerobe. 2005;11(5):247–251. doi:10.1016/j.anaerobe.2005.05.001

19. Wu L, Gingery M, Abebe M, et al. Diversity-generating retroelements: Natural variation, classification and evolution inferred from a large-scale genomic survey. Nucleic Acids Res. 2018;46(1):11–24. doi:10.1093/nar/gkx1150

20. Liu M, Deora R, Doulatov SR, et al. Reverse transcriptase-mediated tropism switching in Bordetella bacteriophage. Science (80-). 2002;295(5562):2091–2094. doi:10.1126/science.1067467

21. Doulatov S, Hodes A, Dal L, et al. Tropism switching in Bordetella bacteriophage defines a family of diversity-generating retroelements. Nature. 2004;431(7007):476–481. doi:10.1038/nature02833

22. Krishnamurthy SR, Wang D. Origins and challenges of viral dark matter. Virus Res. 2017;239:136–142. doi:10.1016/j.virusres.2017.02.002

23. Bianciotto V, Bondi C, Minerdi D, Sironi M, Tichy H V., Bonfante P. An obligately endosymbiotic mycorrhizal fungus itself harbors obligately intracellular bacteria. Chemtracts. 1998;11(3):206–211. doi:10.1128/aem.62.8.3005-3010.1996

24. Cole JR, Wang Q, Fish JA, et al. Ribosomal Database Project: Data and tools for high throughput rRNA analysis. Nucleic Acids Res. 2014;42(D1):633–642. doi:10.1093/nar/gkt1244

25. Wingett SW, Andrews S. Fastq screen: A tool for multi-genome mapping and quality control [version 1; referees: 3 approved, 1 approved with reservations]. F1000Research. 2018;7(0):1–12. doi:10.12688/f1000research.15931.1

26. Bolger AM, Lohse M, Usadel B. Trimmomatic: A flexible trimmer for Illumina sequence data. Bioinformatics. 2014;30(15):2114–2120. doi:10.1093/bioinformatics/btu170

27. Magoč T, Salzberg SL. FLASH: Fast length adjustment of short reads to improve genome assemblies. Bioinformatics. 2011;27(21):2957–2963. doi:10.1093/bioinformatics/btr507

28. Bankevich A, Nurk S, Antipov D, et al. SPAdes: A new genome assembly algorithm and its applications to single-cell sequencing. J Comput Biol. 2012;19(5):455–477. doi:10.1089/cmb.2012.0021

29. Gurevich A, Saveliev V, Vyahhi N, Tesler G. QUAST: Quality assessment tool for genome assemblies. Bioinformatics. 2013;29(8):1072–1075. doi:10.1093/bioinformatics/btt086

30. Seemann T. Prokka: rapid prokaryotic genome annotation. Bioinformatics. 2014;30(14):2068–2069. doi:10.1093/bioinformatics/btu153

31. Finn RD, Bateman A, Clements J, et al. Pfam: The protein families database. Nucleic Acids Res. 2014;42(D1):222–230. doi:10.1093/nar/gkt1223

32. Kelley LA, Mezulis S, Yates CM, Wass MN, Sternberg MJE. The Phyre2 web portal for protein modeling, prediction and analysis. Nat Protoc. 2015;10(6):845–858. doi:10.1038/nprot.2015.053

33. Almagro Armenteros JJ, Tsirigos KD, Sønderby CK, et al. SignalP 5.0 improves signal peptide predictions using deep neural networks. Nat Biotechnol. 2019;37(4):420–423. doi:10.1038/s41587-019-0036-z

34. Lopes A, Tavares P, Petit MA, Guérois R, Zinn-Justin S. Automated classification of tailed bacteriophages according to their neck organization. BMC Genomics. 2014;15(1):1–17. doi:10.1186/1471-2164-15-1027

35. Sharifi F, Ye Y. MyDGR: A server for identification and characterization of diversity-generating retroelements. Nucleic Acids Res. 2019;47(W1):W289–W294. doi:10.1093/nar/gkz329

36. McNair K, Bailey BA, Edwards RA. PHACTS, a computational approach to classifying the lifestyle of phages. Bioinformatics. 2012;28(5):614–618. doi:10.1093/bioinformatics/bts014

37. Bolduc B, Jang H Bin, Doulcier G, You Z, Roux S, Sullivan MB. vConTACT?: an iVirus tool to classify double-stranded DNA viruses that infect Archaea and Bacteria. 2017:1–26. doi:10.7717/peerj.3243

38. Larkin MA, Blackshields G, Brown NP, et al. Clustal W and Clustal X version 2.0. Bioinformatics. 2007;23(21):2947–2948. doi:10.1093/bioinformatics/btm404

39. Gouy M, Guindon S, Gascuel O. SeaView version 4: A multiplatform graphical user interface for sequence alignment and phylogenetic tree building. Mol Biol Evol. 2010;27(2):221–224. doi:10.1093/molbev/msp259

40. Carver T, Berriman M, Tivey A, et al. Artemis and ACT: viewing, annotating and comparing sequences stored in a relational database. Bioinformatics. 2008;24(23):2672–2676. doi:10.1093/bioinformatics/btn529

41. Sullivan MJ, Petty NK, Beatson SA. Easyfig: a genome comparison visualizer. Bioinformatics. 2011;27(7):1009–1010. doi:10.1093/bioinformatics/btr039

42. Bushnell B, Rood J, Singer E. BBMerge – Accurate paired shotgun read merging via overlap. PLoS One. 2017;12(10):e0185056. https://doi.org/10.1371/journal.pone.0185056.

43. Li H, Handsaker B, Wysoker A, et al. The Sequence Alignment/Map format and SAMtools. Bioinformatics. 2009;25(16):2078–2079. doi:10.1093/bioinformatics/btp352

44. Lorenz Christian Reimer, Anna Vetcininova, Joaquim Sardà Carbasse Carola Söhngen, Dorothea Gleim, Christian Ebeling, Jörg Overmann, BacDive in 2019: bacterial phenotypic data for High-throughput biodiversity analysis, Nucleic Acids Research, Volume 47, Issue D1, 08 January 2019, Pages D631–D636, https://doi.org/10.1093/nar/gky879; Bacteroides uniformis. Definitions. 2020. doi:10.32388/9nem1h

45. Leplae R, Lima-Mendez G, Toussaint A. ACLAME: A CLAssification of mobile genetic elements, update 2010. Nucleic Acids Res. 2009;38(SUPPL.1):57–61. doi:10.1093/nar/gkp938

46. Belcaid M, Bergeron A, Poisson G. The evolution of the tape measure protein: Units, duplications and losses. BMC Bioinformatics. 2011;12(SUPPL. 9):S10. doi:10.1186/1471-2105-12-S9-S10

47. Weinberg Z, Lünse CE, Corbino KA, et al. Detection of 224 candidate structured RNAs by Comparative analysis of specific subsets of intergenic regions. Nucleic Acids Res. 2017;45(18):10811–10823. doi:10.1093/nar/gkx699

48. Arambula D, Wong W, Medhekar BA, et al. Surface display of a massively variable lipoprotein by a legionella diversity-generating retroelement. Proc Natl Acad Sci U S A. 2013;110(20):8212–8217. doi:10.1073/pnas.1301366110

49. Xu Q, Shoji M, Shibata S, et al. A Distinct Type of Pilus from the Human Microbiome. Cell. 2016;165(3):690–703. doi:10.1016/j.cell.2016.03.016

50. Ye Y. Identification of diversity-generating retroelements in human microbiomes. Int J Mol Sci. 2014;15(8):14234–14246. doi:10.3390/ijms150814234

51. Goodacre N, Aljanahi A, Nandakumar S, Mikailov M, Khan AS. A Reference Viral Database (RVDB) To Enhance Bioinformatics Analysis of High-Throughput Sequencing for Novel Virus Detection. mSphere. 2018;3(2):1–18. doi:10.1128/mspheredirect.00069-18

52. Gregory AC, Zablocki O, Howell A, Bolduc B, Sullivan MB. The human gut virome database. bioRxiv. January 2019:655910. doi:10.1101/655910

53. Rampelli S, Turroni S, Schnorr SL, et al. Characterization of the human DNA gut virome across populations with different subsistence strategies and geographical origin. Environ Microbiol. 2017;19(11):4728–4735. doi:10.1111/1462-2920.13938

54. Yinda CK, Vanhulle E, Conceição-Neto N, et al. Gut Virome Analysis of Cameroonians Reveals High Diversity of Enteric Viruses, Including Potential Interspecies Transmitted Viruses. Roossinck MJ, ed. mSphere. 2019;4(1):e00585–18. doi:10.1128/mSphere.00585-18

55. Den Besten G, Van Eunen K, Groen AK, Venema K, Reijngoud DJ, Bakker BM. The role of short-chain fatty acids in the interplay between diet, gut microbiota, and host energy metabolism. J Lipid Res. 2013;54(9):2325–2340. doi:10.1194/jlr.R036012

56. Wu GD, Chen J, Hoffmann C, et al. Linking long-term dietary patterns with gut microbial enterotypes. Science (80-). 2011;334(6052):105–108. doi:10.1126/science.1208344

57. Arumugam M, Raes J, Pelletier E, et al. Enterotypes of the human gut microbiome. Nature. 2011;473(7346):174–180. doi:10.1038/nature09944

58. Arumugam M, Raes J, Pelletier E, et al. Enterotypes in the landscape of gut microbial community composition. Nature. 2013;3(1):1–12. doi:10.1038/nature09944.Enterotypes

59. Mahnic A, Id MR. Different host factors are associated with patterns in bacterial and fungal gut microbiota in Slovenian healthy cohort. 2018:1–17.

60. Qin J, Li R, Raes J, et al. A human gut microbial gene catalogue established by metagenomic sequencing. Nature. 2010;464(7285):59–65. doi:10.1038/nature08821

61. Cieplak T, Soffer N, Sulakvelidze A, Sandris D. A bacteriophage cocktail targeting Escherichia coli reduces E. coli in simulated gut conditions, while preserving a non-targeted representative commensal normal microbiota. Gut Microbes. 2018;0(0):1–9. doi:10.1080/19490976.2018.1447291

62. Dalmasso M, Strain R, Neve H, et al. Three new Escherichia coli phages from the human gut show promising potential for phage therapy. PLoS One. 2016;11(6):1–16. doi:10.1371/journal.pone.0156773

63. Krylov V, Shaburova O, Krylov S, Pleteneva E. A genetic approach to the development of new therapeutic phages to fight Pseudomonas aeruginosa in wound infections. Viruses. 2012;5(1):15–53. doi:10.3390/v5010015

64. Mills S, Hill C, Coffey A. Movers and shakers?: Influence of bacteriophages in shaping the mammalian gut microbiota Movers and shakers Infl uence of bacteriophages in shaping the mammalian gut microbiota. 2013;(December 2015). doi:10.4161/gutm.22371

65. Tetz G V., Ruggles K V., Zhou H, Heguy A, Tsirigos A, Tetz V. Bacteriophages as potential new mammalian pathogens. Sci Rep. 2017;7(1):1–9. doi:10.1038/s41598-017-07278-6

